# SECRET-GWAS: Confidential Computing for Population-Scale GWAS

**DOI:** 10.1101/2024.04.24.590989

**Authors:** Jonah Rosenblum, Juechu Dong, Satish Narayanasamy

## Abstract

Genomic data from a single institution lacks global diversity representation, especially for rare variants and diseases. Confidential computing can enable collaborative GWAS without compromising privacy or accuracy, however, due to limited secure memory space and performance overheads previous solutions fail to support widely used regression methods. We present SECRET-GWAS: a rapid, privacy-preserving, population-scale, collaborative GWAS tool. We discuss several system optimizations, including streaming, batching, data parallelization, and reducing trusted hardware overheads to efficiently scale linear and logistic regression to over a thousand processor cores on an Intel SGX-based cloud platform. In addition, we protect SECRET-GWAS against several hardware side-channel attacks, including Spectre, using data-oblivious code transformations and optimized speculative load hardening. SECRET-GWAS is an open-source tool and works with the widely used Hail genomic analysis framework. Our experiments on Azure’s Confidential Computing platform demonstrate that SECRET-GWAS enables multivariate linear and logistic regression GWAS queries on population-scale datasets (one million patients, four million SNPs, 12 covariates) from ten independent sources in just 4.5 and 29 minutes, respectively.

---

Genome-wide association studies (GWAS) correlate genetic variants to phenotypes (traits, diseases) [1]. They help us understand genetic risks for diseases and develop targeted drugs [2]. GWAS are conducted using a genomic database, which typically consists of a set of patient’s genetic variants (millions of SNPs) along with phenotypes, ancestry information, environmental factors, etc. that serve as covariates.

Many institutions have started sequencing their patients’ DNA as sequencing technology gets faster and cheaper to use [3]. For example, the Michigan Genomic Initiative (MGI) has sequenced over 91,000 patients and integrated their data with electronic health records [4]. MGI is currently one of the largest single-institution genomic databases. Even so, its genomic data is not large and diverse enough to represent the global population, especially for rare genetic variants and rare diseases. It is also well known that many racial groups are significantly under-represented in genomic and health datasets [5].

Multiple institutions collaborating allow us to perform population-scale GWAS with millions of patients, which can address issues of population diversity and rare variant/disease representation. We need to pool sensitive genomic data from different institutions to derive statistically powerful insights through GWAS. Pooling data also increases available sample sizes for most racial groups, rare genetic variants, and rare diseases.

However, governmental (e.g. GDPR) and institutional regulations require protections for patient privacy. Unfortunately, we currently lack a computing system solution that can pool data from multiple sources and perform population-scale GWAS, while also preserving individual patient’s privacy.

Today, when health data is shared (e.g., MCI’s Genomic data commons [4]), privacy is regulated with a cumbersome approval process and user agreements. Unfortunately, institutions lack mechanisms to enforce privacy agreements and ensure that approved users are not misusing the data. Another common approach is to perform local analyses and share only summary statistics which are then combined in a meta-analysis [6]. However, this greatly reduces the fidelity of results and therefore is not a viable solution [6, 7]. These low-confidence privacy solutions inhibit institutions from sharing data.

Previous studies [7–13] use homomorphic encryption (HE) and/or cryptographic multi-party computation (MPC) to enable privacy-preserving collaborative GWAS. These methods are several orders of magnitude slower than plaintext computation [14], meaning GWAS with population-scale datasets may take *many months or even years* which is computationally infeasible. HE is a form of encryption that allows computation on encrypted data without revealing any information about the plain text. MPC is a technique that enables independent parties to exchange non-sensitive values to jointly compute on the sensitive data without revealing anything private. The most efficient related work is Froelicher et. al [7], a HE/MPC hybrid solution that requires over four days to complete a query on a fairly small dataset (20k patients, 4 million SNPs, and 12 covariates) using 72 processor cores. Such an analysis with millions of patients would optimistically take many months but depending on the number of SNPs and covariates could take years.

Confidential computing uses hardware to guarantee privacy, however, challenges in limited hardware resources have prevented population-scale GWAS using common statistical analyses. Intel’s propriety implementation of hardware enclaves known as Software Guard eXtensions (SGX) provides strong security guarantees with significantly lower overheads compared to HE and MPC [14]. Institutions send encrypted data to a central SGX enclave node. SGX ensures data confidentiality by guaranteeing that data remains encrypted in main memory during computation. Data is converted to plaintext only within the trusted processor. All external parties, including the operating system, are prevented from observing secret values in the user’s trusted enclave program. SGX also has cryptographic checks that guarantee the integrity and freshness of the data and computation, a feature that HE/MPC methods lack. Thus, SGX guarantees the privacy of computation even on an untrusted third-party system such as a public cloud, where multiple institutions can pool sensitive data to perform collaborative GWAS as depicted in Figure 1.

**Fig. 1:**
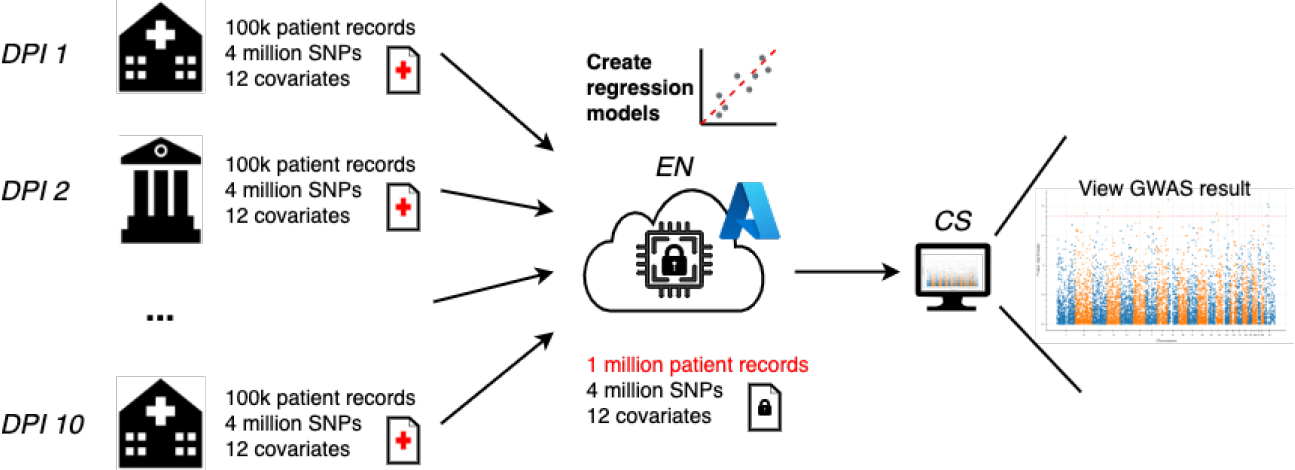
Computational nodes in collaborative SECRET-GWAS. DPI nodes send encrypted data to Enclave Nodes (EN) on a confidential computing cloud. ENs perform privacy-preserving GWAS. A coordination server (CS) orchestrates computation and collates output from ENs.

While prior works have explored using SGX for privacy-preserving genome analysis, they are limited to simpler statistical analyses such as TDT [14], simplified PCA, and chi-squared association tests [15]. It has not yet been shown that SGX can support widely used linear and logistic regression GWAS kernels. This is due to the limited secure enclave memory available in SGX (at most 256 MB) which is too small to hold genomic datasets that range from many GBs to several TB in size. Furthermore, collaborative GWAS using linear and logistic regression with population-scale datasets (*≥*1 million patients) has never been demonstrated with *any* privacy-enhancing technology (HE, MPC, confidential computing, etc.)

We present the <u>S</u>calable <u>E</u>fficient <u>C</u>onfidential and privacy-<u>R</u>especting <u>E</u>nvironment for <u>T</u>rusted GWAS (SECRET-GWAS), an SGX-based confidential computing solution for population-scale collaborative GWAS. **Through several systems and SGX-specific performance optimizations, we show for the first time that we can perform linear and logistic regression GWAS queries for datasets with a million patients in a few minutes**. We also show that our massively parallel implementation scales to over 1000 cores on Microsoft’s Azure confidential computing platform.

SECRET-GWAS consists of three primary optimizations: streaming with data locality, batching, and data parallelism. Each SNP can be processed independently of others. Therefore, we only need to store the data required to process a single SNP within the enclave memory at any given time. We also observe that a significant fraction of a SNP’s working set (patient’s covariates) is a constant across all SNPs, and it fits well within the small enclave memory.

Our second optimization significantly reduces the SGX overheads by reducing the frequency of function calls into and out of enclaves. We achieve this by batching as many SNPs together as possible while ensuring that the working set does not overflow the enclave memory. Lastly, testing for genetic association of a given SNP does not require information about any other SNPs, and therefore computation of each analysis is independent. We leverage this insight to build a parallel system capable of scaling to any number of processor cores. We present a solution that evenly distributes SNP partitions across over a thousand cores in a synchronization-free manner and ensures that the work is load-balanced across the partitions.

We also protect SECRET-GWAS from side-channel attacks against the trusted hardware through two defensive techniques. Our first defense is a data-oblivious implementation of linear and logistic regression kernels. It is well understood that conditional branches and memory accesses dependent on sensitive data can leak private information [16–21]. We address this problem by using oblivious code transformations where all sensitive conditional branches are predicated, ensuring that the control flow and memory access sequence are independent of sensitive data input. In other words, our oblivious GWAS kernel will always execute the same sequence of instructions and memory accesses regardless of the input. This eliminates the timing-based and memory access pattern side-channel attacks.

Our second defense is against Spectre [22], which can leak information due to speculative optimizations in hardware such as branch prediction. We defend against Spectre by applying an LLVM compiler pass designed to harden loads against speculative attacks [23, 24]. We optimize this further by selectively using predicated execution for branches where the predication is faster than the Spectre mitigation.

SECRET-GWAS is open-source and hosted on GitHub (https://github.com/jonahrosenblum/SECRET-GWAS). Our experiments were all deployed on Azure’s confidential computing cloud platform. SECRET-GWAS supports both linear and logistic regression and is integrated with the widely used genomic analysis platform Hail [25], which is often used for analysis in peer-reviewed GWAS papers. Hail handles data quality control and filtering in the GWAS pipeline, and our system produces genetic association outputs consistent with Hail (which has no privacy protections). SECRET-GWAS is capable of performing real-world collaborative GWAS, as it can scale to population-scale datasets used in genomic studies [7].

Our experiments focus on evaluating the performance of the SECRET-GWAS system, which is independent of the data being synthetic or authentic. We conduct an SGX-based GWAS that pools data from 10 different source nodes and analyzes it using over 1000 processor cores in the cloud. All source nodes are geographically distributed over a wide area network spanning from the East to the West Coast of the United States. Our GWAS uses a dataset with 4 million SNPs and 12 covariates per patient. We query with both 20,000 and 1 million patients, finding that GWAS with linear regression takes 10 seconds and 4.5 minutes, respectively, while GWAS with logistic regression is completed in 45 seconds and 29 minutes, respectively.

## Results

To fully understand our threat model and gain some helpful background information please see our Background and Motivation section.

### Overview of SECRET-GWAS

In a standard collaborative GWAS pipeline, there are multiple independent institutions with their own private health and genomic databases. Each institution is responsible for performing quality control, filtering, and preprocessing of their dataset (which can be done offline). These processed records are then pooled together and analyzed SNP by SNP, performing association tests on each one while controlling for factors such as patient ancestry, age, sex, etc. The final result is a file listing every SNP and the association likelihood with the disease being studied.

The goal of SECRET-GWAS is to enable the genetic association step of a GWAS for mutually untrusting institutions without leaking any private patient data, which has not been done before at a population-scale with popular association tests such as regression. We achieve this by using hardware enclaves to protect the sensitive computation of linear and logistic regression models on Azure’s confidential computing cloud platform.

As shown in Figure 1, SECRET-GWAS consists of the following entities:

#### Data Providing Institution (DPI)

Each institution is assumed to have its own private health and genomic database. DPIs are responsible for quality control, filtering, and preprocessing their data into the Hail file format. They partition their dataset evenly by SNPs, then encrypt and send each partition to the appropriate enclave node.

#### Enclave Node (EN)

This is an SGX-enabled cloud instance (Microsoft Azure in our study) that receives encrypted patient records for a given SNP from all the sources (DPIs). It builds linear and logistic regression models for each SNP within the privacy-preserving SGX enclave. The non-sensitive output is then sent to the GWAS organizer.

#### Coordination Server (CS)

This centralized server orchestrates the communication between DPIs and ENs. It also aggregates the GWAS results generated by the ENs.

Only the secure enclave in the EN is trusted in our threat model. Our pipeline assumes each DPI handles its quality control, filtering, and preprocessing offline using Hail [25]. Patient records are pooled in the ENs and all sensitive computation occurs inside the secure hardware enclaves. The CS’s only pipeline-related role is acting as an aggregator that compiles the final result from outputs generated by ENs.

Many GWAS pipelines perform PCA to correct for population stratification. While SECRET-GWAS is capable of using population PCs to control for ancestry, computing them collaboratively with PCA is not part of our current pipeline. Additionally, while we assume the results of genetic association tests (i.e. test statistics) are non-sensitive, some prior works have added noise to these results via differential privacy [26, 27]. Future works may address both limitations by adding privacy-respecting collaborative PCA and differential privacy to the SECRET-GWAS pipeline.

### Streaming to Overcome Enclave Memory Size Limitations

When a program runs inside a hardware enclave, the secure region of protected memory that it can use is called the Enclave Page Cache (EPC). As noted in our Challenges section, the EPC size in main memory is relatively small (128-256 MB [28]) compared to the standard main memory size (dozens of GBs). Datasets used in GWAS are typically hundreds of GB or even TBs in size, depending on the number of patients and SNPs. Therefore, it is crucial to design a GWAS system such that the working data set can fit within the small EPC to avoid prohibitive EPC paging overheads.

We can build a regression model for a SNP independent of others, as there is no data dependency between their computations. We refer to this as *SNP-level parallelism*. To compute the regression model for a SNP, we need the covariates (*c*) and the SNP’s genotype value for all patients (*p*). Thus, the working data set size for processing a SNP is on the order of *p ∗* (*c* + 1).

If we use 8 bytes per covariate value, then even with 1 million patients and 12 covariates we only require about 100 MB of main memory, which fits well within the EPC (128-256 MB in size).

We also observe that covariate records (about 100 MB in the previous example) are constant for all SNPs. We exploit this temporal data locality by copying them only once into the EPC and reusing them across all SNPs computed on an enclave node.

Thus, to process a new SNP, we only need to copy its genotype values for all patients into the EPC and discard them after building that SNP’s regression model. This data is a small fraction 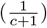 of a SNP’s working set.

### Batching to Reduce Enclave Boundary Transfers

To enter or exit a secure enclave program, special function calls, ECalls and OCalls respectively, must be used. As we describe in our Challenges section, each ECall and OCall incur significant performance overhead because they trigger a context switch and TLB flush. If we process each SNP independently, then we need to make these calls for every SNP to stream their data (refer to our section on Streaming) in and out of the enclave memory.

To solve this problem, we group several SNPs into a batch, so that a boundary call is required only once per batch. There are diminishing returns for batch size - if the batch is too large, then the working set will exceed the EPC, incurring EPC paging overhead. Therefore, we maximize the batch size under the constraint that its working set size fits within EPC. We calculate the optimal batch size in terms of SNPs to be 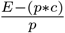 where *E* is the size of the EPC, *p* is the number of patients, and *c* is the number of covariates.

### Data Parallelism

Harnessing modern processors’ parallel computing capabilities has driven significant performance gains across a myriad of applications. As mentioned in our Challenges section, SNPs can be analyzed independently of others, which we call *SNP-level parallelism*. The number of SNPs is typically the largest dimension in a GWAS dataset (several million). While this parallelism has been noted in literature [7, 9, 13], there are no trusted hardware-based privacy-preserving GWAS systems [7, 15] that exploit this parallelism across multiple SGX machines.

A key challenge in parallelization is ensuring that computation is evenly distributed among computational nodes. Otherwise, the node with the largest partition and therefore the most computation would become the bottleneck. In the case of collaborative GWAS, this imbalance arises due to the fact not all the institutions sequence the same SNPs (refer to our Implementation section for more details). Without coordination between DPIs, it is not guaranteed that SNPs will be evenly and correctly partitioned across ENs.

We address this problem locally at the source node (DPI) while partitioning the SNPs. We use a hashing scheme [29] that creates a deterministic but pseudorandom mapping between SNPs and partitions. Although it is pseudorandom, this hashing scheme we use has been shown to deviate from the ideal distribution with a standard error of 7.64E-9 [29]. This ensures that on average each partition will be processed in the same amount of time.

### Side-channel Mitigation

One of the concerns for creating applications with trusted hardware like SGX is side-channel attacks [17, 18, 30], where attackers can extract sensitive data by measuring the architectural state during program execution. Therefore, in SECRET-GWAS we develop novel data oblivious GWAS kernels and apply an optimized speculative load hardening to defend against important side-channels such as control flow [16, 17], memory access [18–21], and Spectre [22]. We go into more detail about side-channels in our Threat Model and Challenges sections. Further, details of our mitigations for floating-point timing side-channels are in our Implementation and Artifacts section.

#### Data Oblivious Kernel

The control flow of a program and the memory access patterns may depend on sensitive data which can leak information from an enclave [16–18, 31, 32]. Because different branch paths perform different computations, measuring the execution length of a program can oftentimes reveal information about the data used in the conditional branch. For example, a popular algorithm used in regression for GWAS is calculating the determinant of a matrix. A common variant of this algorithm does a memory swap operation only if a sensitive value is zero. By measuring the execution time of this algorithm, an attacker can infer the number of zeros in the input. Alternatively, an attacker could monitor memory accesses (e.g., by snooping the memory bus transactions) to determine whether a memory swap is being done or not, and from that infer the number of zeros in the sensitive input data. We defend against these side-channels by implementing data-oblivious GWAS kernels for linear and logistic regression. These customized kernels are guaranteed to execute the same sequence of instructions and access the same sequence of memory locations irrespective of the sensitive input data. This is accomplished using predicated execution [33]. The idea is to execute both paths of a conditional branch, regardless of the condition. Then, using mathematical operators, only the correct path’s result is stored in the output variables. Figure 2 provides simplified examples of all oblivious transformation patterns we used in our kernels. The code transformations in the figure are simplified for readability. These predicated executions are sufficient to ensure that the memory access sequences are uniform across all sensitive inputs. The sequence length may vary depending on the dataset size, but this does not leak information about individual patient records.

**Fig. 2:**
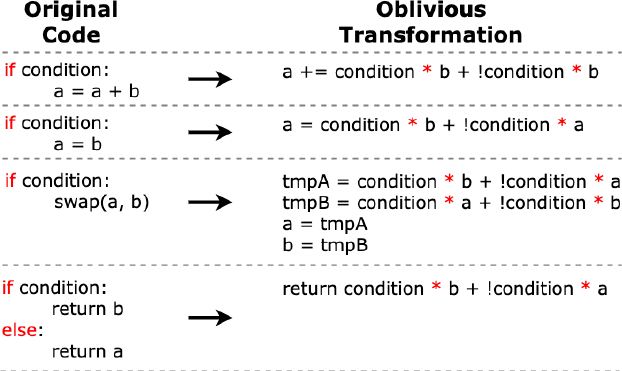
Oblivous transformations used in SECRET-GWAS. Conditional branches are rewritten using predicated execution, as high-lighted in red.

Lastly, we ensure the compiler cannot violate our data oblivious guarantee by modifying our predicated executions to use conditional branches. All predicated executions in our code-base are implemented with handwritten branch-free assembly code that the compiler cannot modify, thus guaranteeing oblivious execution.

#### Efficient Spectre Mitigation

Spectre is a side-channel vulnerability that tricks a victim program into speculatively executing code that changes the architectural state of the machine, and then later measures that change to extract sensitive information [22]. The most popular variant of this involves using branch mispredictions to load sensitive data into the cache and then measuring how long non-speculative loads take to learn the sensitive value in the cache [22]. This attack works even when the control flow does not depend on sensitive data, so our data-oblivious transformations do not protect us from Spectre.

We apply an LLVM compiler pass developed by Google’s Project Zero [23, 24] that defends against the Spectre branch speculation/-cache attack by preventing loads from happening inside of speculated branches until the predcate can be resolved [22, 24]. This defense treats any load that occurs in a conditional branch as potentially dangerous and prevents the load from accessing the memory address until the predicate of the branch has resolved. This hardened load technique stops Spectre attacks from using speculation to alter the state of the cache and therefore prevents data leakage from branch/load attacks.

The Spectre load hardening mitigation introduces a new performance burden. Although our oblivious kernels remove all branches on sensitive data using predication [33], there are still many non-sensitive branches whose performance is impacted by the load hardening mitigation. We observe that predication removes the possibility of Spectre branch/load attacks, as it eliminates speculation. With this in mind, we predicate several non-sensitive conditional branches in our kernels, as long as they have short paths, resulting in a performance improvement. For longer paths, predicated execution is more expensive than Spectre defense.

Most modern processors are shipped with patches for some Spectre vulnerabilities, however there are still trusted processors used today with the vulnerability not fixed. If users are not concerned about Spectre attacks the defense can easily be turned off by changing a single compiler flag.

### Methodology

All of our experiments were conducted on Microsoft Azure, a public cloud platform. Execution times are averaged from at least 3 runs. Experiments for Table 1, Figure 4 and Figure 5 were performed on a WAN, with the DPIs located in the US East datacenter and ENs located in the US West datacenter to conservatively simulate geographical distance between institutions.

**Table 1:**
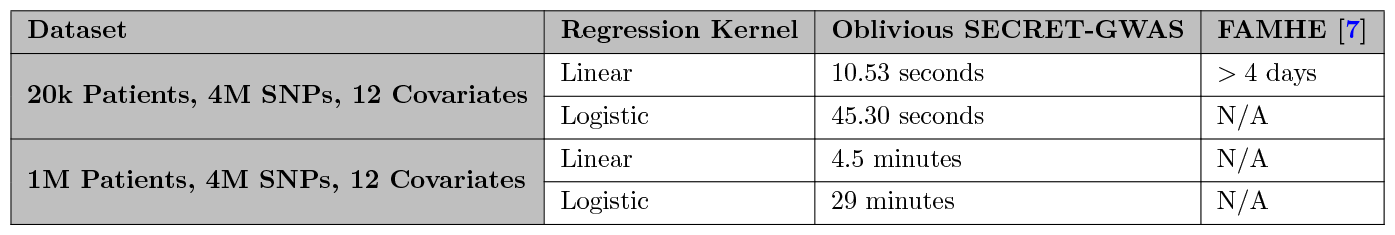
SECRET-GWAS and FAMHE [7] execution times.

All the DPIs and CS (non-SGX machines) run on 3rd Gen Intel Xeon Platinum 8370C processors with 64 cores. ENs (SGX-enabled) run on 3rd Gen Intel Xeon Scalable processors. We use a single core for all experiments, except for experiments that study parallel performance (Table 1, Figure 4, and Figure 5), where many dual-core machines were used concurrently. For the experiment in Figure 3c, we used SGX-enabled Intel Xeon E-2288G processors of size DC2s with 56 MB of EPC and DC4s with 112 MB of EPC to test the effects of using the same processor with two different EPC sizes.

**Fig. 3:**
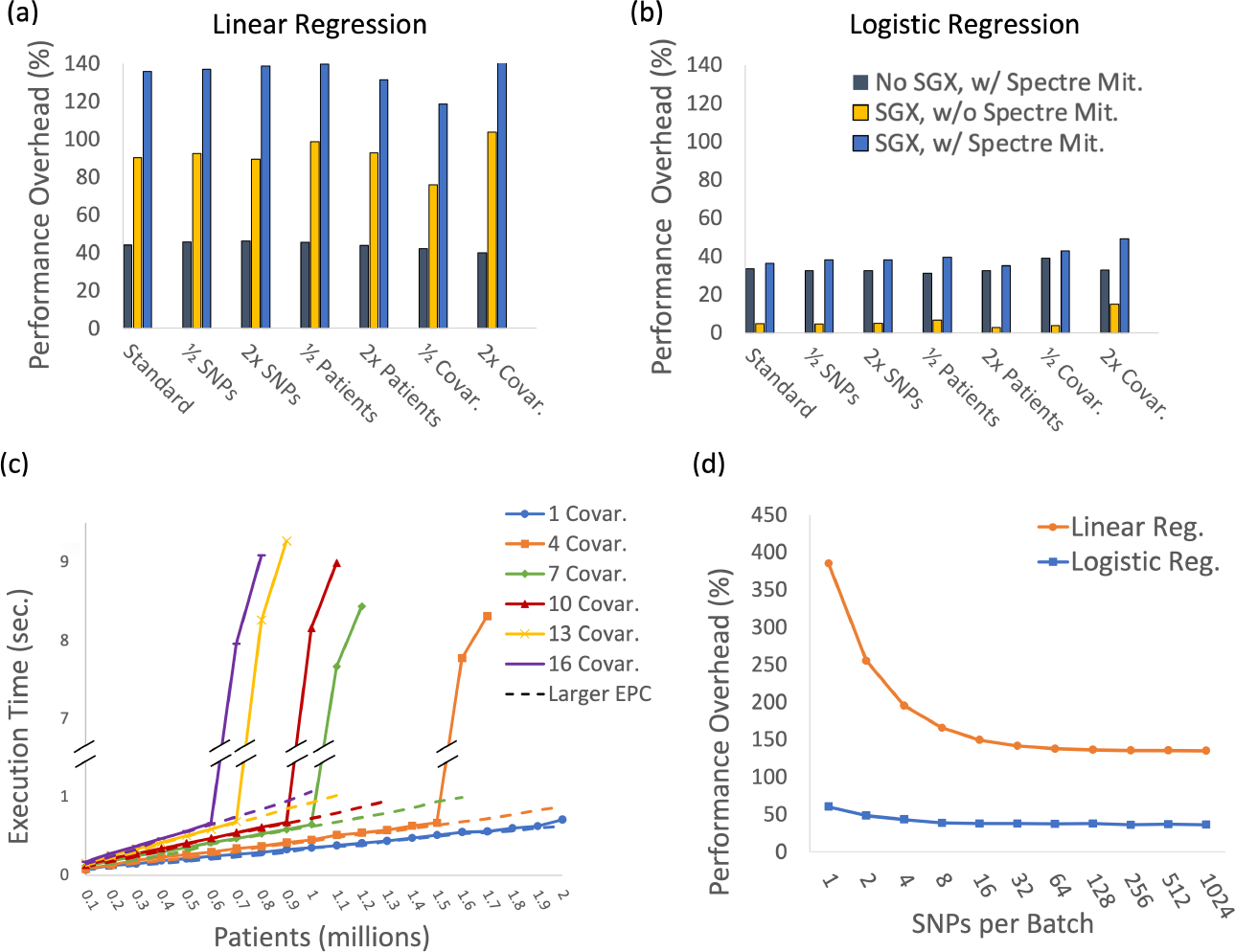
Figures (a) and (b) show SGX performance overhead and sensitivity to dataset size. Figure (c) shows how well SECRET-GWAS scales with the number of patients and covariates used in the GWAS. Additionally, we test EPC sensitivity to show how the performance goes super linear when the 56 MB EPC overflows and paging is triggered, but when the larger 112 MB EPC is swapped in (dotted line) the performance becomes regular again. Figure (d) shows the effectiveness of SNP batching in lowering the performance overhead of both linear and logistic regression.

Additionally, the SGX machines used in Azure’s confidential compute platform have all known SGX vulnerability mitigations implemented with microcode updates and added protections in silicon [17]. All experiments use compilation flags -O3 and -mllvm -x86-speculative-load-hardening except for one portion of Figures 3a and 3b which are labeled as “SGX, w/o Spectre Mit.” All genomic datasets used are upsampled from a chunk of the 1000 Genomes Project dataset [34]. We study several dataset configurations. One is **20K-Data**, similar to FAMHE’s largest dataset [7], which consists of 20k patients, and each patient record contains 12 covariates and 4 million SNPs. Another is **1M-Data**, a large dataset representing population-scale GWAS with a million patients, and the same number of SNPs and covariates as **20k-Data**. Other configurations are explained below along with the experiments that use them.

### GWAS Accuracy

The results generated by SECRET-GWAS were compared to Hail to measure accuracy. Both systems use the same method of imputation to ensure that missing genotypes are handled similarly. Several datasets were used to generate GWAS output files for both Hail and SECRET-GWAS. We tested two Hail kernels: `logistic_regression_rows` using the Wald test and `linear_regression_rows`. Through our testing, we confirmed that the results generated by SECRET-GWAS have the same accuracy as Hail.

### Performance Comparisons

SECRET-GWAS is the first GWAS system to support privacy-preserving linear and logistic regression using SGX. Table 1 shows the execution times on 1024 cores for linear and logistic regression on two different datasets, each of which is spread across 10 different data sources.

SECRET-GWAS can perform GWAS at interactive speed for **20K-Data**: 10.5s and 45.3s for linear and logistic regression, respectively. SECRET-GWAS can also handle **1M-Data** in 4.5min and 29min for linear and logistic regression.

We compare our performance to the fastest known privacy-preserving solution for regression, which is FAMHE [7]. FAMHE is a hybrid MPC/HE GWAS system that is capable of performing GWAS with linear regression on **20K-Data** in *>*4 days using 72 processor cores [7]. MPC performance does not scale linearly with the number of cores, as it is limited by network bandwidth. FAMHE demonstrated that doubling their processor cores used from 72 to 144 improved performance only by 28%.

Froelicher et al. showed that FAMHE’s execution time grows linearly with the number of patients [7]. Based on this, we estimate that it would take over 210 days to perform the GWAS using linear regression with a million patients, which is computationally infeasible. We did not find a privacy-preserving GWAS solution that could perform logistic regression for our dataset sizes.

### Privacy (SGX) Overhead

Figures 3a and 3b show the overheads due to Spectre mitigations and SGX to make SECRET-GWAS privacy-preserving. We do this by studying four system configurations. The first is a baseline version of SECRET-GWAS that uses no SGX or Spectre mitigations. All performance overheads shown are normalized to this version. Second is SECRET-GWAS with Spectre mitigations but without SGX. The third configuration, adds SGX but removes Spectre mitigations. The final configuration is the true unmodified SECRET-GWAS with both Spectre and SGX protections used.

For the “standard” dataset (125k SNPs, 10k patients, 6 covariates), we find that SECRET-GWAS incurs 135% and 36% performance over-head for linear and logistic regression respectively. These overheads are orders of magnitude less compared to HE/MPC methods. We observe interesting differences in the source of overhead between kernels. Both kernels have a similar overhead due to the Spectre mitigation (about 30-40%). But logistic regression has negligible SGX overhead (5%) compared to linear regression (90%). We attribute this to logistic regression being a compute-bound rather than memory-bound kernel, and the SGX overheads are proportional to the rate of cache misses to memory. Logistic regression enjoys higher genotypic data reuse due to its iterative nature and therefore has a lower cache miss rate.

We also study the sensitivity of these over-heads concerning different dimensions of the GWAS dataset. We changed one dimension (SNPs, patients, covariates) at a time by either halving or doubling it; this is labeled as 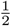 and 2x in Figures 3a and 3b. We find that the number of SNPs does not affect the overhead. While the total amount of work increases with the number of SNPs, computation (and therefore associated overhead) per SNP does not change. We see that overhead reduces marginally for larger datasets with 2x patients. This is because more patients per SNP implies more computation per SNP, which reduces SGX-related overheads as a fraction of overall runtime. The number of covariates has the opposite relationship to that of patients. As noted in our Background section, SGX over-heads are sensitive to cache misses. While both covariate and genotype data fit into the EPC in main memory, they do not fit within a processor’s cache. Therefore, more covariates in the dataset increase the cache miss rate, which in turn results in higher SGX overhead.

### Effectiveness of Batching Optimization

We presented our batching optimization to reduce the frequency of enclave calls (ECalls and OCalls). We tested the effectiveness of our optimization again using the “standard” dataset (125k SNPs, 10k patients, 6 covariates). As seen in Figure 3d, SGX performance overhead decreases significantly with large batches. With 1024 SNPs per batch, the overhead is 134% for linear regression and 36% for logistic regression. Without batching (batch size of 1) the overhead jumps to 385% for linear regression and 60% for logistic regression. We observe more savings for linear regression, as it is more memory-bound than logistic regression. Enclave calls trigger TLB flushes, which cause memory-bound kernel performance to suffer more. By reducing enclave call frequency, batching yields significant performance benefits.

### EPC Memory Scaling

As mentioned in Section 2, we will incur significant performance degradation if the working data set size does not fit within the EPC. We study this problem for GWAS in Figure 3c by using the same processor with two different EPC configurations from Azure. Each run only uses 20 SNPs since thousands of GWAS experiments were run when gathering data (240 different dataset/EPC configurations, and averaging across multiple runs) to create Figure 3c.

Execution time increases linearly with the number of patients until the working set overflows the EPC size (for our first Azure configuration this is 56 MB), at which point the performance goes superlinear. A second dotted line shows that if the EPC is increased to 112 MB, the performance will continue to scale linearly with the number of patients. This experiment demonstrates that the performance degradation is due to EPC paging, and not due to any other factor. In all our experiments for SECRET-GWAS, except for the EPC sensitivity test in Figure 3c, we did not trigger EPC paging. Recent work shows that increasing the scalability of the EPC is possible [35, 36], and Intel has developed new approaches to increase EPC memory in SGX [28] that have recently been added to Azure’s cloud offerings. Therefore, we believe that GWAS will be able to support even larger datasets with several million patients as the pace of EPC technology improves.

### Parallel System Scaling

Figure 4 shows how well our system performance scales with the number of processor cores. We vary the number of cores for a fixed dataset size. Because we partition SNPs evenly across all cores in the system, more cores mean each core processes fewer SNPs. All experiments in Figure 4 used a dataset with 4 million SNPs, 6 covariates, and the labeled number of patients.

**Fig. 4:**
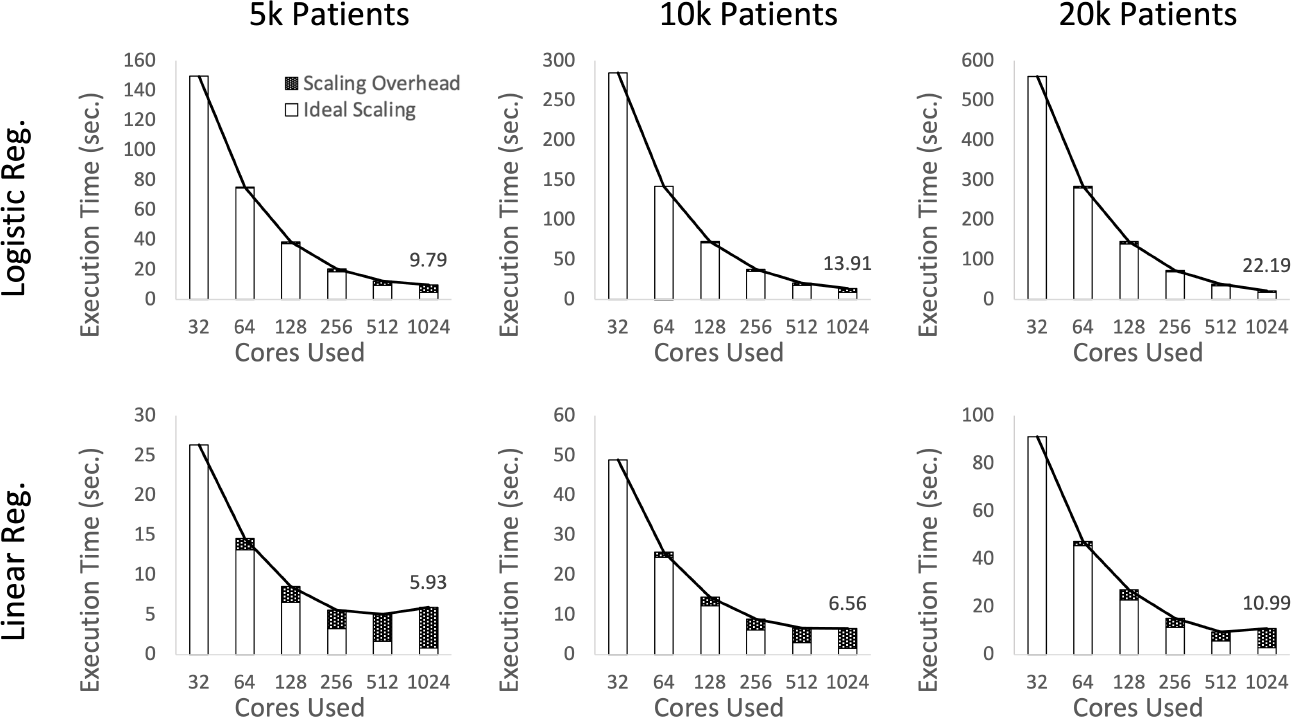
Performance of SECRET-GWAS scales linearly with processor cores.

For logistic regression, we see nearly perfect scaling as we get linear speedup with cores. The same is true for linear regression up to around 512 cores. For datasets with a larger number of patients, we observe better scaling. To further understand the scaling overheads, we compute an ideal speedup for a given number of cores and then attribute the remaining execution time to scaling overhead. We see that scaling overheads are significant only for linear regression at high core counts.

Linear regression has less compute per SNP, and therefore scaling overheads due to network and synchronization overheads start to dominate as the amount of work per core reduces to a negligible amount. Figure 5 provides further evidence where we see that linear regression requires a significantly higher proportion of networking threads compared to logistic regression. Another reason for scaling overhead is that the bottleneck shifts to data sources (DPIs) and CS as we scale the number of cores in the enclave nodes (ENs). Increasing the number of cores and network bandwidth available to the CS and DPIs is a solution to this problem.

**Fig. 5:**
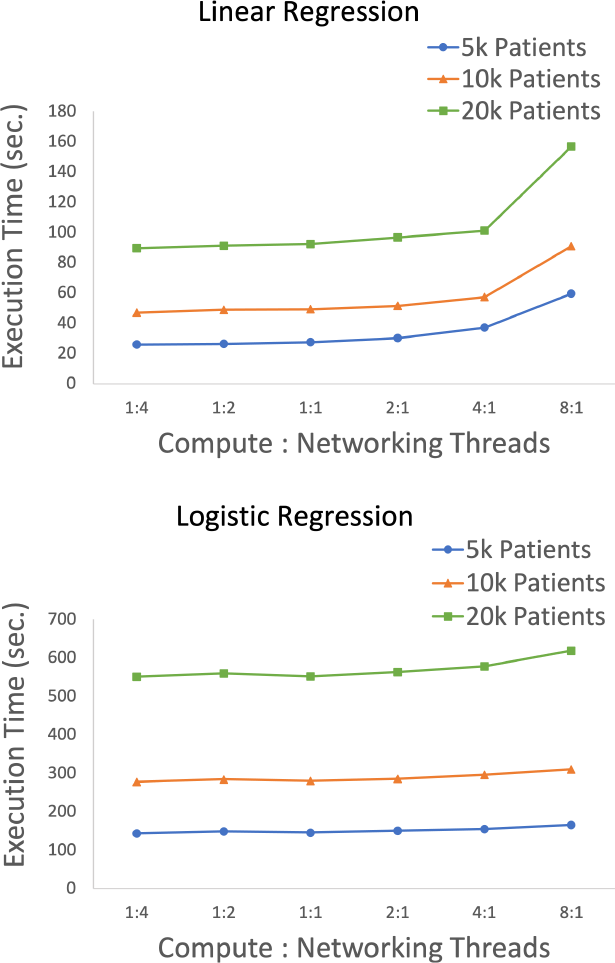
Performance of SECRET-GWAS for different ratios of compute to networking threads.

## Discussion

This paper presents SECRET-GWAS, a confidential computing solution for population-scale GWAS. Previous solutions were limited in the complexity of the kernels and/or datasets they supported. Through a set of system optimizations, SECRET-GWAS enables fast linear and logistic regression analysis over a large dataset with a million patient records, 4 million SNPs, and 12 covariates in 4.5 and 29 minutes respectively. SECRET-GWAS also presents the first oblivious transformations to harden GWAS kernels against side-channel attacks. Additionally, we protect against Spectre and floating-point timing attacks by applying optimized speculative load hardening and disabling computations with subnormal numbers. By deploying SECRET-GWAS on Azure’s confidential computing platform, we show that its performance scales linearly with every dimension of a GWAS dataset, and can parallelize to over processor 1000 cores.

Future works may look to enhance the SECRET-GWAS pipeline by adding more genetic association kernels (e.g. SVM), support for PCA to control for ancestry, and obfuscation of computed statistics via differential privacy.

In conclusion, SECRET-GWAS is a large first step towards secure collaborative GWAS at a population-scale.

## Online Methods

### Background and Motivation

This section first describes our threat model. We then provide a brief background on confidential computing, its advantages, and challenges in applying it to GWAS.

### Threat Model

We seek a privacy-preserving solution where institutions can pool data in a centralized server such as a public cloud service, or an institution’s private data center. All entities in our system are explained in our Design section. Of these, the hardware enclave running on the EN is the *only* trusted entity in our threat model. We do not trust the DPIs, CS, the cloud provider, the study coordinator, etc. and they do not need to trust each other.

Our security goal is to protect patient privacy and avoid sensitive data leaks. Denial of service attacks and tampering with the study results are *not* part of our threat model. We assume that the cloud service provider’s middleware software such as their operating system, their system administrators, and those with physical access to the machine are not trusted. For our adversary, denial-of-service attacks are trivial (i.e. unplugging the machine). Additionally, while we guarantee that the computation happening *inside* the enclave will be correct, we do not prevent the untrusted DPIs from contributing bogus data to the GWAS. The DPIs can spoil the GWAS result by sharing fake patient records, but can not compromise the privacy of any patient record held by other institutions. Additionally, we do not trust the CS that organizes communication between parties and aggregates results - this untrusted entity never views or influences the handling of sensitive information.

Our threat model includes common dangerous side-channel attacks that take advantage of program differences stemming from conditional branching and memory access patterns, as well as Spectre attacks. We are also aware of a less common side-channel attack that measures timing differences in floating point operations [17, 37]. This is also part of our threat model, and more details on how we address it can be found at the end of our Implementation and Artifacts section. Analog side-channels that measure the power draw and heat of the processor during execution require expensive equipment to measure and are incredibly difficult to pull off. As a result, they are not included in our threat model.

We assume that the publicly available GWAS software used to analyze sensitive genomics data is trusted, in that it guarantees all its outputs (file and network) do not contain sensitive information. We also assume that the GWAS output is non-sensitive and can be publicly shared with all participating institutions. Some prior works [26, 27] challenge this assumption and further obfuscate GWAS outputs via differential privacy, but this is orthogonal to the problem we solve in our paper. Additionally, there is no privacy-preserving technology (including HE and MPC) that solves this problem implicitly, meaning it is not unique to trusted hardware or our work.

Finally, we assume that the trusted processor used to execute our GWAS software (Intel SGX [38] in our implementation) is correct and bug-free.

### Confidential Computing

Confidential computing (CC) uses processor hardware (e.g., Intel SGX [38]) as root-of-trust to create a trusted execution environment (TEE) called “enclaves”. Application developers define enclave functions that can operate on sensitive data, allocate enclave memory, and copy sensitive data in and out of the enclave. CC on an untrusted system (e.g., public cloud) guarantees that the enclave’s code and data are protected from the rest of the system. All other software, including the operating system, and those with physical access to the system will not be able to tamper with or observe the enclave’s execution, code, or data. More specifically, Intel SGX guarantees the following security properties:

#### Confidentiality

All enclave memory is encrypted while outside the processor (main memory, storage). Enclave memory is only decrypted within the trusted processor where it is inaccessible by any adversary [38].

#### Integrity

Message authenticating hashes are appended to all enclave memory proving it has not been tampered with. The hashes are stored outside the trusted processor. If tampering is detected the execution is aborted [38].

#### Freshness

Only the most up-to-date values are kept in enclave memory. If a malicious user tries to “replay” an old value SGX will detect it via the on-die Merkle tree [38].

#### Attestation

Users can verify with Intel that the SGX device they are using is properly configured and is running the code that they expect [38]. It is worth noting that HE does not guarantee integrity without also using computationally expensive Zero-Knowledge Proofs [39]. Additionally, both HE and MPC do not provide freshness or attestation guarantees. No privacy-preserving technology is strictly better than another.

### Challenges

There are several performance challenges in using SGX for GWAS.

#### Small Enclave Memory Size

The size of enclave memory, or enclave page cache (EPC), that supports freshness is small by modern standards. A larger EPC would grow the size of the Merkle tree and increase the already high cache-miss performance overhead (19.5% and 102% for writes and reads respectively) making it prohibitively expensive [40]. SGX, for instance, only provides 128-256 MB of EPC [28]. If an application’s working data set size exceeds EPC size, then it triggers EPC paging - an expensive process where encrypted pages are moved back and forth between non-enclave and enclave memory, slowing down programs by 5x on average [35]. Datasets used in population-scale GWAS can be several GBs, which do not directly fit within the EPC. Previous studies on SGX for GWAS used compression and sampling methods to partially overcome this problem and yet were still limited to simpler statistical analysis such as TDT and Pearson’s chi-squared test, while tasks like PCA required approximation [14, 15]. We address this problem by exploiting data locality in GWAS to enable linear and logistic regression that was not previously shown to be possible.

#### Enclave Call Overheads

SGX supports function calls into and out of the trusted enclave using entry calls (ECalls) and out calls (OCall) [40]. Both functions incur a significant performance overhead because they trigger context switches and TLB flushes [38]. We present optimizations that reduce the frequency of these calls.

#### Parallelism

Lastly, previous works on SGX for GWAS have not demonstrated scaling of parallel execution past a single SGX server using 8 cores [15]. Our solution automatically works with any number of machines, cores, and threads. We exploit the high degree of SNP-level parallelism in GWAS by scaling SECRET-GWAS to over a thousand processor cores.

#### Side-Channel Vulnerabilities

The applications running inside of trusted hardware such as SGX may be vulnerable to side-channel attacks [18, 31, 32]. Side-channels are a class of security attacks that observe the physical behavior of an application, and based on this infer sensitive information. For example, suppose a secure application takes three times as long to process records containing rare diseases. In that case, an attacker can measure how long each record takes to process to determine which ones have rare diseases. This type of attack is concerning when we consider a strong threat model, where an attacker controls the operating system software and has physical access.

Popular side-channel attacks discussed in prior works such as control flow timing [16, 17], memory access pattern analysis [18–21], as well as Spectre attacks [22] are also part of our threat model. To address this we use a novel combination of oblivious transformations and speculative load hardening to defend against the aforementioned side-channels.

### Implementation and Artifacts

Our SGX application uses Microsoft’s OpenEnclave library [41], an open-source SGX SDK. OpenEnclave was chosen due to its portability and multi-platform support. It also contains many useful cryptography libraries.

SECRET-GWAS is integrated with Hail, a popular Python genomic analysis framework. Hail is designed to run at scale in a cloud environment and is easily customized by researchers through the Hail open-source Python library. SECRET-GWAS uses Hail for quality control and filtering capabilities, and can read in Hail format files. The resulting output files closely mimic Hail.

We have implemented a demo to show how Hail is integrated with the SECRET-GWAS ecosystem. We provide two Python scripts in our demo - one uses Hail for all functionality, and one uses SECRET-GWAS for performing regression analysis. The two output files can be easily compared to see the similarity. Lastly, our imputation method is the same as Hail’s and imputes using the average value for a given SNP.

SECRET-GWAS is designed to automatically scale to any number of cores. To configure the system, the user just needs to specify the number of ENs, the number of DPIs, and the IP address of the CS in easily modifiable JSON files - SECRET-GWAS figures out the rest. We also provide scripts written for the Azure CLI to quickly spin up hundreds of machines equipped with SGX, as well as scripts for cleaning up Azure resources after the GWAS concludes.

To scale to over a thousand independent cores, several implementation details were key. For example, using AES-CBC with 256-bit keys is important to protect confidentiality and authenticity while data is in transit over the network. Using this standard, however, means that cores cannot share keys without considerable synchronization overhead. We instead issue every core (*C*) one AES key per DPI (*D*) in the system, meaning we have *C ∗ D* unique AES keys in use for the duration of a GWAS. The keys are only communicated once the DPIs attest and verify that the CNs are running the correct enclave and are configured properly.

Additionally, to reduce network latency and improve efficiency overall we use the PLINK format for storing the information on one genotype in 2 bits [42]. We considered using efficient encodings and SNP compression techniques, which can save memory and network resources[14, 15, 43]. We decided not to, as standard compression is vulnerable to side-channel privacy leaks as the performance of these tools and memory access patterns are input data dependent.

Lastly, we have identified that GWAS rarely, if ever, utilizes the subnormal range of floating point numbers. This is especially true when double-precision floating points are used for computations, something done in both SECRET-GWAS and Hail [25]. Therefore, we opt to disable subnormal numbers by setting both the flush-to-zero and denormals-to-zero flags using the MXCSR state management instructions. Prior works have shown that setting these flags in SSE processors fixes the performance discrepancy and therefore the side channel attack [37]. If setting these flags fails we throw an error to the user that their machine may be vulnerable to this floating point side-channel attack.

### Related Work

We discuss previous work on using trusted hardware for GWAS, MPC/HE-based solutions, and other applications that use SGX.

#### Trusted Hardware for GWAS

PRINCESS [14] supports transmission disequilibrium tests (TDTs) for 5 million SNPs. The majority of the computation happens at the DPI, with the institution summing up transmitted and untransmitted allele counts for each SNP. These counts are aggregated in an SGX-enabled centralized node to perform TDT. Thus, the individual patient records need not be sent to the central node in PRINCESS. However, for linear and logistic regression models, to enable fast and accurate computation we communicate encrypted records to the enclave node. The working data set size for TDT is also much smaller than what we need to support regression models. PRINCESS uses lossless compression (range encoding) to reduce network communication, and the data is decompressed within an enclave. Compression is prone to side-channel attacks as we discussed in Section 2, and so we opt for the PLINK encoding scheme [42], which is data oblivious.

SkSES [15] performs Pearson’s association test on 5.5 million SNPs and 128K patients. They use PCA methods (simplified EIGENSTRAT) to control for ancestry. To mitigate enclave memory space constraints, they use lossy sketching (a form of sampling) and Huffman encoding. We do not employ any lossy compression or sampling methods to overcome space constraints, while still supporting larger datasets (a million patients). Also, we support linear and logistic regression kernels, which require more computer and memory space than Pearson’s association test. Finally, none of the previous work on SGX for GWAS used oblivious transformations to mitigate side-channel attacks.

#### MPC and HE

As we discussed in Section 2, MPC and HE solutions such as Froelicher et al. [7]’s work incur orders of magnitude slowdown. In contrast, our solution using trusted hardware incurs no more than 2.3x overhead.

Blatt et al. [9] implements an approximation of logistic regression using HE. They test their system on 16k SNPs, 15k patients, and 3 covariates on a single core which takes 66 minutes for logistic regression. By comparison, our system processes 4 million SNPs, 1 million patients, and 12 covariates (which is 71,666x larger) on 1024 cores in 29 minutes as seen in Table 1. Additionally, we do not use any approximations in our logistic regression giving us higher confidence in our results. Smajlović et al. [12] developed Sequre, a framework for quick development of MPC programs. They implement a full GWAS pipeline and estimate a runtime of 3 weeks to process one million patients with linear regression as the genetic association test. SECRET-GWAS can perform the genetic association step of the pipeline with one million patients using linear regression in 4.5 minutes, as seen in Table 1.

#### Other Applications using SGX

Dokmai et al. [17] created SMac, a privacy-preserving genotype imputation application using SGX. Researchers often have missing genotypes in their datasets and wish to impute them for their analysis. However, doing this imputation is highly sensitive as the data being sent to the imputation server and the data residing on the server are both highly sensitive. SMac solves this problem by using SGX as a trusted platform for imputation. Furthermore, SMac addresses side-channel leakages by fixing sensitive data-dependent timing differences and memory access patterns in their imputation code with code transformations that improve resiliency to side-channels.

Zheng et al. [18] created Opaque, a secure SGX-based distributed analytics platform. Opaque is a platform built on Spark SQL that can perform queries on sensitive data in a secret manner using SGX. To address memory access pattern leaks, Opaque implemented data-oblivious query operations, such as “oblivious sort”. Ohrimenko et al. [30] used SGX to support privacy-preserving machine learning (ML) kernels (support vector machines, neural networks, k-means clustering). ML is similar to GWAS in that both benefit from multiple institutions pooling their sensitive data. To address side-channel due to data-dependent access patterns, Ohrimenko et al. created several data oblivious primitives. These primitives were used to implement oblivious ML algorithms.

Although there are differences in each of these applications, a common concern is addressing side channels through oblivious code transformations. All three aforementioned works found that they had memory access pattern leakages in their applications, and applied either an oblivious transformation [18, 30] or made their subroutines leakage-resistant [17]. Our work is the first solution that transforms GWAS kernels to be data oblivious.

